# First genome of *Labyrinthula*, an opportunistic seagrass pathogen, reveals novel insight into marine protist phylogeny, ecology and CAZyme cell-wall degradation

**DOI:** 10.1101/2020.09.14.297390

**Authors:** Mun Hua Tan, Stella Loke, Laurence J. Croft, Frank H. Gleason, Lene Lange, Bo Pilgaard, Stacey M. Trevathan-Tackett

**Affiliations:** Centre of Integrative Ecology, School of Life and Environmental Sciences Deakin University, Geelong, Australia; Deakin Genomics Centre, Deakin University, Geelong, Australia; School of BioSciences, Bio21 Institute, University of Melbourne, Australia; Department of Microbiology and Immunology, University of Melbourne, Bio21 Institute, Melbourne, Australia; School of Life and Environmental Sciences, University of Sydney, Sydney, 2006, New South Wales, Australia; BioEconomy, Research & Advisory, Valby, 2500 Copenhagen, Denmark; Protein Chemistry and Enzyme Technology, Department of Bioengineering, Technical University of Denmark, Kgs. Lyngby, Denmark

**Author notes:** **Corresponding Author**: Dr Stacey Trevathan-Tackett, Deakin University, 221 Burwood Hwy, Burwood, Victoria, Australia 3125.

**Keywords:** evolution, ion regulation, mitochondrial genome, saprobe, Stramenopiles, virulence

## Abstract

*Labyrinthula* spp. are saprobic, marine protists that also act as opportunistic pathogens and are the causative agents of seagrass wasting disease (SWD). Despite the threat of local- and large-scale SWD outbreaks, there are currently gaps in our understanding of the drivers of SWD, particularly surrounding *Labyrinthula* virulence and ecology. Given these uncertainties, we investigated *Labyrinthula* from a novel genomic perspective by presenting the first draft genome and predicted proteome of a pathogenic isolate of *Labyrinthula* SR_Ha_C, generated from a hybrid assembly of Nanopore and Illumina sequences. Phylogenetic and cross-phyla comparisons revealed insights into the evolutionary history of Stramenopiles. Genome annotation showed evidence of glideosome-type machinery and an apicoplast protein typically found in protist pathogens and parasites. Proteins involved in *Labyrinthula*’s actin-myosin mode of transport, as well as carbohydrate degradation were also prevalent. Further, CAZyme functional predictions revealed a repertoire of enzymes involved in breakdown of cell-wall and carbohydrate storage compounds common to seagrasses. The relatively low number of CAZymes annotated from the genome of *Labyrinthula* SR_Ha_C compared to other Labyrinthulea species may reflect the conservative annotation parameters, a specialised substrate affinity and the scarcity of characterised protist enzymes. Inherently, there is high probability for finding both unique and novel enzymes from *Labyrinthula* spp. This study provides resources for further exploration of *Labyrinthula* ecology and evolution, and will hopefully be the catalyst for new hypothesis-driven SWD research revealing more details of molecular interactions between *Labyrinthula* species and its host substrate.

## Introduction

Sequencing capabilities of both short and long read technologies have shown marked progress in recent years, resulting in the generation of a large number of contiguous and high quality genome assemblies reported for a myriad of living organisms at an unprecedented pace. As a result of genome sequencing projects at local-to-global scales, we are better resolving linages across the Tree of Life (Lewin et al. 2018), revealing new diversity for underrepresented or unculturable microorganisms (e.g. aquatic and marine protists and zoosporic fungi; Sibbald and Archibald 2017, Lange et al. 2019a) and informing the ecological and functional roles of organisms in the environment (Kyrpides et al. 2014). In marine ecosystems, genomic and transcriptomic sequencing is producing vital information on the ecological roles microorganisms play. For example, genomes of putative pathogens are revealing genes involved in virulence and attachment to the host, as well as mechanisms involved in degrading the host substrate and switching between non-pathogenic and pathogenic lifestyles (Fernandes et al. 2011). These new insights are especially important as marine emergent diseases and opportunistic pathogens are predicted to increase globally in the coming decades due to the suboptimal living conditions for marine biota caused by human- and climate-induced changes (Harvell et al. 1999).

Seagrass meadows are one such marine ecosystem susceptible to a decline in health and increased infection, as they are subjected to dynamic pressures at the land-sea interface, such as elevated temperatures in shallow regions and run-off-associated turbidity, eutrophication and hyposalinity conditions (Sullivan et al. 2018). There are currently four pathogens that cause disease to multiple seagrass genera: *Labyrinthula, Phytophthora, Halophytophthora*, and Phytomyxea (Sullivan et al. 2018). *Labyrinthula*, a heterotrophic, halophytic protist in the Stramenopile lineage, is the causative agent of seagrass wasting disease (SWD). To-date, it is also the most well-studied seagrass pathogen after being discovered in the early twentieth century when it decimated 90% of the North Atlantic population of the seagrass *Zostera marina* (Sullivan et al. 2018). *Labyrinthula* isolates from seagrass leaves have shown variable levels of virulence through laboratory infection trials and associated phylogenetic analyses (Martin et al. 2016, Brakel et al. 2017). Virulent isolates are able to invade the leaf cells of living plants and subsequently cause black leaf lesions, the diagnostic trait of SWD (Sullivan et al. 2018). However, *Labyrinthula*, is also a ubiquitous commensal saprobe that decomposes marine plants and algae litter, with one species parasitic to turfgrass (Martin et al. 2016). Because of the flexibility of its ecological function, *Labyrinthula* spp. are considered opportunistic pathogens that in some cases utilise a mild parasitism strategy (Brakel et al. 2017).

Since its discovery, the advances in our understanding of *Labyrinthula* species include phylogenetic reclassifications of *Labyrinthula* (Leander and Porter 2001) and molecular techniques that facilitate detection of *Labyrinthula* without the need for culturing (Duffin et al. 2020, Lohan et al. 2020). Despite this progress, we still do not completely understand the pathosystem of SWD. For example, it is still under debate whether the conditions that promote SWD rely more on seagrass health and defence capabilities or the virulence and enzymatic capabilities of *Labyrinthula* isolates and haplotypes, or a combination of host and pathogen characteristics (Martin et al. 2016, Duffin et al. 2020). Much of the new knowledge on the pathosystem has come from the immune and stress response of the seagrass using genomic markers or assays (Brakel et al. 2014, Duffin et al. 2020), as opposed to pathogen virulence (Martin et al. 2016). Furthermore, despite our molecular advances, we also know little about the role of specific organelles, such as the bothrosome, in the hunt and consumption of food sources (Collier and Rest 2019). Next-generation sequencing technologies are helping us produce new hypotheses for *Labyrinthula* and SWD. For example, Lohan et al. (2020) questioned how the variation in *Labyrinthula* diversity links with intra-species diversity of the seagrass host.

This set of ecological uncertainties constitutes the basis for our interest in sequencing the genome of a *Labyrinthula* isolate. So far the genomes of only three different genera of the Labyrinthulea class have been sequenced, which has been valuable in revealing novel ecological insights and biotechnology applications (Song et al. 2018). In this study, using a hybrid assembly approach combining both short and long read sequence data, we assemble the first known draft genome of a pathogenic haplotype of *Labyrinthula* and report a highly contiguous assembly. Through sequencing and annotating a draft *Labyrinthula* genome, our aim is to further resolve the genus’ lineage and function, particularly its capacity to degrade seagrass carbohydrates, and compare those functions in closely-related but non-pathogenic, saprobic members of Labyrinthulea. We expect that this work will also provide a launching pad to deeply investigate and understand the basic ecology and evolutionary history of this ubiquitous marine protist, as well as to unlock the potential of their understudied and unexploited CAZyme diversity.

## Materials and methods

### *Labyrinthula sp*. isolate description, sample preparation and sequencing

The *Labyrinthula* sp. (order Labyrinthulida) isolate used for sequencing was cultured on seawater-serum agar plates (Trevathan-Tackett et al. 2018) at room temperature prior to preparing DNA and RNA for sequencing. Isolated from a seagrass meadow in Victoria, Australia, isolate SR_Ha_C (GenBank: MF872134.1; referred simply as *Labyrinthula* SR_Ha_C herein) was previously shown to be highly pathogenic to both *Zostera muelleri* and *Heterozostera nigricaulis* (Trevathan-Tackett et al. 2018). Genomic DNA for short-read Illumina sequencing was extracted from the *Labyrinthula* SR_Ha_C culture with the ZymoBIOMICS Quick DNA kit according to the manufacturers’ instructions. Total RNA was prepared using the Monarch Total RNA isolation kit according to the standard protocol for hard to lyse cells. The short-read genome library was prepared with the Nextera Flex DNA library preparation kit (Illumina, San Diego) and sequenced on the MiSeq with 2 x 300bp V3 chemistry. The short-read RNAseq library was prepared with the Nugen mRNA ovation kit and sequenced on the MiniSeq with 2×150bp chemistry. The full-length transcriptome libraries were prepared for long-read sequencing with Lexogen’s teloprime kit and Oxford Nanopore’s SQK-LSK109 library prep kit (ONT, UK). Sequencing was performed on one Flongle (FLO) flow cell for 24 h and one MinION R9.4.1 flow cell for 24 h. See Electronic Supplementary Material 1 for detailed isolate, extraction, assembly and annotation methodology.

### Genome assembly

Illumina reads were trimmed for quality and adapters with Trimmomatic, whereas adapters were trimmed from Nanopore MinION reads with Porechop. Haploid genome size, repetitive content and heterozygosity size for *Labyrinthula* SR_Ha_C was estimated with GenomeScope based on the frequency of k-mers in short reads (Vurture et al. 2017). MaSuRCA (Zimin et al. 2017) was used to perform a hybrid assembly of the genome, combining short and long read data. Resulting contigs were corrected and polished with three iterations of Racon and Pilon. Since bimodal k-mer distributions were observed from the GenomeScope analysis, indicating high levels of heterozygosity, the Purge Haplotigs pipeline was used to remove haplotigs (Roach et al. 2018). Checks were put into place to scan for possible bacterial contaminant contigs. The first was to search for any contigs with first hits to bacterial sequences in NCBI’s non-redundant nucleotide database. Secondly, RNAmmer was used to identify contigs containing bacterial ribosomal RNAs. In both searches, none of the contigs met these criteria and therefore all were retained. The final assembly was compared to those of other Stramenopiles based on assembly quality and completeness with Quast (Mikheenko et al. 2018) and BUSCO (Seppey et al. 2019), respectively. A list of species included in this comparison is available in Table S1.

### Mitogenome recovery and phylogenetic analysis

The complete *Labyrinthula* SR_Ha_C mitogenome was extracted from the assembly through *blastn* homology with other protist mitogenomes. The mitogenome was annotated with MFannot and MITOS and manually adjusted according to proteins in NCBI’s non-redundant database. Phylogenetic reconstruction was performed based on thirteen genes typically found in eukaryote mitogenomes *(atp6, atp8, cox1-3, nad1-6, nad4l, cytb)* (Boore 1999). These were extracted from 127 Stramenopile mitogenomes and four from outgroup species (Table S2). Protein sequences were aligned with MAFFT, trimmed with trimAl and concatenated into a superalignment with FASconCAT-G. IQ-TREE was used to find the best-fit partition model and to infer a maximum-likelihood tree. The tree was rooted with *Reclinomonas americana*, which has a mitogenome that most closely resembles the ancestral proto-mitochondrial genome (Lang et al. 1997).

### Gene prediction and functional annotation

Two types of “hints” were used to predict genes for this non-model organism. The first type consists of *Labyrinthula* transcripts generated in this study by sequencing (Illumina) and assembling the transcriptome (Haas et al. 2013) to generate an Illumina-based transcriptome (Data S1), in addition to long Nanopore-sequenced cDNA reads. The second type included known peptide sequences from 235 protists downloaded from Ensembl Protists (release 46; Howe et al. 2019) and curated sequences from UniProtKB/Swiss-Prot (Consortium 2018).

Gene prediction was performed with the MAKER2 annotation pipeline (Holt and Yandell 2011). A first iteration was run to identify a preliminary set of genes based on provided hints. The predictions were used to train two *ab initio* gene predictors, Augustus and SNAP, after which the resulting gene models were used in a second MAKER2 iteration. Since protist genomes are diverse and therefore contribute highly divergent gene sequences, the *keep_preds* option was activated in this iteration to report all predictions regardless of concordance to supporting hints. InterProScan was used to scan these predictions for the presence PFAM protein domains and any gene models that have physical evidence (annotation edit distance, AED < 1) and/or protein domains were retained. To annotate function, DIAMOND (Buchfink et al. 2015) was used to find homology between predicted genes and proteins in NCBI’s non-redundant protein database (e-value cutoff 1e-5).

Ortholog discovery within the kingdom Stramenopila (i.e. Chromista) was performed using protein sequences obtained from GenBank for two commonly-studied protists: 1. *Plasmodium falciparum* (BioProject: PRJNA148), 2. *Phytophthora sojae* (BioProject: PRJNA262907). Pairwise searches of protein sequence sets were performed with DIAMOND (Buchfink et al. 2015) and orthologs between *Labyrinthula* and the two species were identified through Reciprocal Best Hits (RBH).

### Identification and peptide-based functional annotation of putative carbohydrate-active enzymes

The dbCAN2 meta server (Zhang et al. 2018) was used to annotate carbohydrate-active enzymes (CAZymes) based on enzyme families available in the CAZy database (Cantarel et al. 2008). Due to the divergence of *Labyrinthula* from other eukaryotes, the HMMER search option was selected (E-Value < 1e-15, coverage > 0.35) to search against representative hidden Markov models of signature domains within the five enzyme classes: Glycoside Hydrolases (GH), Glycosyl Transferases (GT), Polysaccharide Lyases (PL), Carbohydrate Esterases (CE), and Auxiliary Activities (AA). Counts of putative *Labyrinthula* SR_Ha_C CAZyme sequences were compared to data available on the CAZy database for 36 other Stramenopile species.

The predicted amino acid sequences of *Labyrinthula* SR_Ha_C were analysed by the new non-alignment based method, Conserved Unique Peptide Patterns (CUPP; Barrett and Lange 2019). CUPP automatically, annotates proteins to families and subfamilies. Further, if characterised members of the CUPP group query is already assigned to a specific CAZyme function, CUPP assigns that function to the query protein. For the purpose of comparison, predicted proteins of three other members of the class Labyrinthulea (a.k.a. Labyrinthulomycetes) were analysed: *Aplanochytrium kerguelense* (PBS07; family Aplanochytridae, order Labyrinthulida), as well as *Aurantiochytrium limacinum* (ATCC MYA-1381) and *Schizochytrium aggregatum* (ATCC 28209) (family Thraustochytriidae, order Thraustochytrida).

## Results & Discussion

### A contiguous *Labyrinthula* sp. genome

This study reports the first assembled genome of a *Labyrinthula* species, available on GenBank (BioProject: PRJNA607370, WGS version: JAALGZ01). This 50.9 Mbp assembly contained 159 contigs (Table 1), consistent with the kmer-based method’s estimate of a 51 Mbp haploid genome size for this isolate (Figure S1). The majority of short reads (99.07%) were successfully aligned back to the assembly. Contiguity and completeness of this assembly were substantially improved by the inclusion of Nanopore long reads in a hybrid assembly, demonstrating the value of strategic incorporation of reads from different platforms. Though the SR_Ha_C isolate was maintained in culture media that included streptomycin and penicillin, further careful checks were performed to exclude bacterial contigs. No bacterial contaminants were found in the genome, ensuring that findings reported in this study accurately reflect the genetics of this *Labyrinthula* SR_Ha_C isolate. The ‘clean’ genome will also prevent inaccurate interpretations in other larger-scale studies looking into investigations of bacteria-to-animal lateral gene transfer (Sibbald and Archibald 2017).

**Table 1.** Data generation, genome assembly and annotation of the *Labyrinthula* sp. isolate SR_Ha_C. *nr* refers to the non-redundant protein database. aa = amino acids.

|  |  |
| --- | --- |
| **Raw sequencing data and depth** (based on 51 Mbp genome size estimate) |  |
| Illumina (MiSeq) | 21.5 Gbp (421.79×) |
| Oxford Nanopore (MinION) | 612 Mbp (12.00×) |
| **Mitogenome assembly** |  |
| Assembly size (circular) | 26,822 bp |
| # protein coding genes | 23 |
| # ribosomal RNAs | 2 |
| # transfer RNAs | 36 |
| **Genome assembly** |  |
| Number of contigs | 159 |
| Assembly size | 50,948,119 bp |
| Contig N<sub>50</sub> (median) size | 1,265,572 bp |
| Average contig size | 320,428.42 bp |
| Longest contig | 3,302,845 bp |
| Shortest contig | 4,362 bp |
| **BUSCO completeness (lineage: *stramenopiles_oddb10*, n:100)** |  |
| Complete | 60.0% |
| Fragmented | 9.0% |
| Missing | 31.0% |
| **BUSCO completeness (lineage: *eukaryota_oddb10*, n:255)** |  |
| Complete | 42.8% |
| Fragmented | 9.0% |
| Missing | 48.2% |
| **Genome annotation** |  |
| Number of ab initio proteins predicted by gene predictors | 17,041 |
| with AED<sup>+</sup> < 1 and/or with PFAM domain | 8,488 |
| with homology to NCBI's *nr* database (1e-5) | 6,351 |
| with homology to NCBI's *nr* database (1e-50) | 1,372 |
| Average protein length | 470 aa |
| Average number of exons/gene | 2.03 |
| Longest protein* | 15,723 aa |

<sup>+</sup> Annotation edit distance (AED) ranges from 0 to 1, a small value shows lesser variance between the predicted gene and provided evidence.
\* Homologous to “Receptor-type tyrosine-protein phosphatase F” [*Hondaea fermentalgiana*]

Genome completeness was assessed with the BUSCO algorithm, which reported a 60.0% and 42.8% completeness for this reported assembly, based on *stramenopiles_odb10* and *eukaryota_odb10* ortholog sets, respectively (Table 1, Figure 1). The former ortholog set (27 species, 100 BUSCOs) contained only three species from the phylum Bigyra compared to the Oomycota (14) and Ochrophyta (9) phyla. The latter (70 species, 255 BUSCOs) was derived from data of 37 metazoan species, 22 fungi, 10 plants and one Rhodophyta, the single protist representative in this dataset. While the *Labyrinthula* SR_Ha_C assembly reported relatively high values of longest contig and N50 lengths, BUSCO-assessed completeness ranked lower than that of most other Stramenopiles (Figures 1a, S2). Higher completeness was generally observed for species in the phylum Oomycota but appeared to be more variable in the other two phyla (Figure 1a). In contrast, QUAST analysis showed *Labyrinthula* SR_Ha_C outperforming the Bigyra phylum average (green cells in Figure 1b) across all measured metrics (Figure 1c). Similarly, high quality QUAST outcomes for *Blastocystis hominis* were not reflected in BUSCO completeness values (*Labyrinthula* SR_Ha_C 60%, *Blastocystis hominis* 45%).

**Figure 1:**
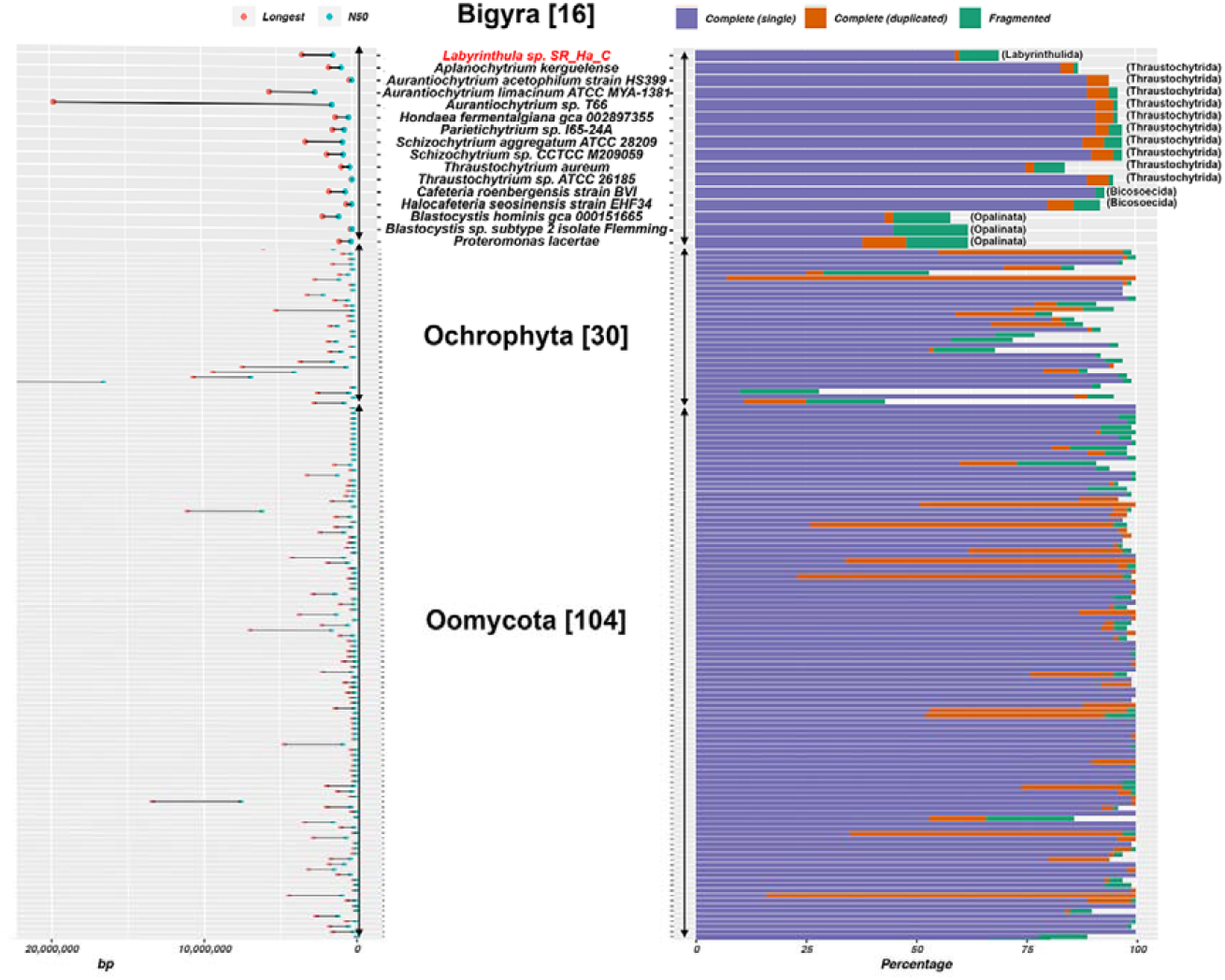
Comparison of genome assemblies for Stramenopiles in three phyla: Bigyra, Ochrophyta and Oomycota. a. (Left) N_50_ length and longest contig/scaffold for each assembly (cutoff 20 Mbp); (Right) Percentages of complete and fragmented BUSCOs based on 100 BUSCOs in *stramenopiles_odb10*. Number of assemblies per phylum are listed in brackets. A detailed version and sources are available in Figure S2 and Table S1. b. Further details on assemblies with sequence (contig/scaffold) length distribution for species in the phylum Bigyra. Longest sequence length, N_50_ and N_75_ are converted to percentages out of assembly length for more accurate comparisons across species with different genome sizes. Green cells are for values that outperform the average and red for values poorer than average, subject to the context of each column. c. QUAST-generated Figures showing a cumulative length plot, Nx values and GC content for species in the phylum Bigyra (colour legend in panel b). Assembly sources: Ensembl Protists, NCBI, JGI.

Though a surge of new genomes are being made available in more recent times, taxonomic sampling of protists are largely skewed to select ‘popular’ lineages with relevance to medical, agricultural and aquaculture interests. However, since gene model prediction, quality assessment and other comparative genomic analyses largely rely on the support of closely-related sequences; this patchy and uneven representation presents gaps in current knowledge and challenges efforts to better characterise and understand genomes of less-studied organisms. In the case of quality assessments, while the BUSCO algorithm has been shown to be an effective and popular choice of tool for assessing the ‘completeness’ of genome assemblies, its shortcomings have also been discussed especially in relation to lineages sampled in its reference ortholog datasets (Expositor□Alonso et al. 2020). Subject to characteristics of an assessed genome, there can be technical limitations in estimating proportions of fragmented and missing BUSCOs, and in species that are highly divergent from the reference ortholog dataset, higher degrees of reported BUSCO ‘missingness’ may reflect the organism’s evolutionary divergence and history rather than the completeness of its assembly (Exposito□Alonso et al. 2020). Our results show that the use of multiple assessment qualities is important to more accurately assess the quality of assemblies, particularly those of highly divergent, non-model organisms (Exposito□Alonso et al. 2020).

### *Labyrinthula* sp. gene annotation

A transcriptome with 1.2 million transcripts was generated and assisted in the annotation of the genome, resulting in the prediction of >17,000 protein sequences (Table 1; annotation results in Data S1). We applied a conservative filter to retain only gene models that were either supported by existing sequence evidence or those that contain PFAM protein domains, in order to avoid false gene predictions albeit at the cost of removing some real, but clade-specific, novel genes. Since the annotation process typically relies on the homology of sequences to those in other well-annotated reference genomes (Exposito□Alonso et al. 2020), the absence of a genome available as a close relative of *Labyrinthula* SR_Ha_C resulted in a reduced final number of 8,488 proteins reported in this study (Table 1; Data S1). This is likely an underestimation of the true number owing to the limited sequenced genomes in this clade and further highlights the novelty of the *Labyrinthula* genome.

A survey of the annotated functions of these predicted peptides revealed rhodanese-like proteins related to stress response and sulphur metabolism (Cipollone et al. 2007), as well as a rhodopsin protein (Data S1). While light-activated functions have not been explored in *Labyrinthula* species, phototaxis and rhodopsin-related functions have been increasingly identified for estuarine zoosporic fungi and protists (Kazama 1972, Tian et al. 2018). We noted proteins involved in sodium, potassium and calcium exchange and transport suggestive of the pathways involved in ion homeostasis and salinity adaptations in halophytic organisms (Harding et al. 2017). The annotation also revealed a wealth of actin-myosin and carbohydrate-degradation proteins. A pairwise annotation, with orthologs of known pathogenic protist *Plasmodium falciparum* and oomycete *Phytophthora sojae*, further revealed glideosome and apicoplast ribosomal proteins (Data S1). The presence of a glideosome protein in addition to the actin-myosin machinery suggests shared traits with the apicomplexan family of pathogens, and opens an interesting avenue for investigating the machinery underlying *Labyrinthula* parasitism and host invasion, as well as its evolutionary history of endosymbiosis (Keeley and Soldati 2004, Oborník et al. 2009).

### CAZymes and functional predictions facilitating plant cell wall degradation

A total of 112 genes associated with carbohydrate-active enzyme (CAZy) activities were identified from the predicted *Labyrinthula* gene set (Figure 2, Table S3). This was distributed across 56 glycosyltransferases (GT), 34 glycoside hydrolases (GH), 17 auxiliary activities (AA), 3 polysaccharide lyases (PL) and 2 carbohydrate esterases (CE). We identified the presence of genes categorised in several groups of putative aromatic and cellulose-degrading enzymes with activities associated with the breakdown of plant cell walls. Some of these enzyme families also show an enrichment of genes relative to other available protist data on the CAZy database, including, the AA2 enzyme family (activities: manganese peroxidase, lignin peroxidase), the PL1 family (activities: pectate lyase, pectin lyase), the GH1 family (activities: ß-glucosidases, ß-galactosidases) and the GH5 family (activities: endoglucanase/cellulase, etc) (Figure 2).

**Figure 2:**
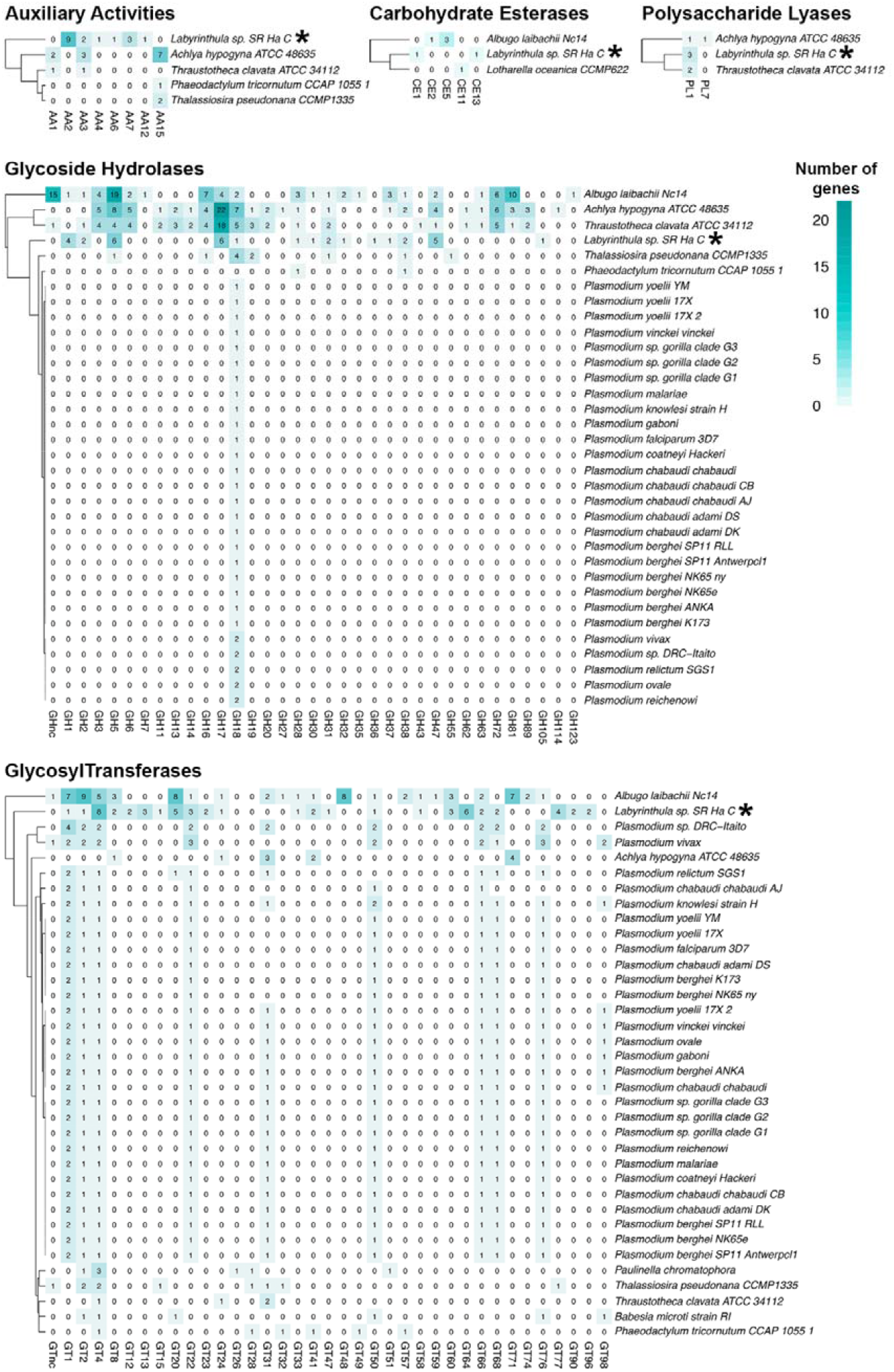
Annotation of Carbohydrate-active enzymes in *Labyrinthula* sp. isolate SR_Ha_C in comparison with other Stramenopile profiles available in the CAZy database. Comparative analysis was done for five enzyme classes: Auxiliary Activities (AA), Carbohydrate Esterases (CE), Glycoside Hydrolases (GH), GlycosylTransferases (GT) and Polysaccharide Lyases (PL). Enzyme family numbers correspond to those in the CAZy database and are indicated on the x-axis. Asterisks (*) indicate the *Labyrinthula* isolate SR_Ha_C reported in this study.

As these enzyme families can act on a diversity of substrates, we used the CUPP analysis to more closely predict the function of the CAZymes. We found that the *Labyrinthula* SR_Ha_C genome has a broad selection of cell wall-degrading and storage materials modifying enzymes (starch, cellulose, trehalose, Beta-D-galactosides, α-mannan. α-1,2-mannan), as well as glucosylceramidase that can be involved in pathways in plant hosts-pathogen relationships (Table 2) (Warnecke and Heinz 2003). The three enzyme functions with highest number of functionally annotated genes were 1,2-α-mannosidase (5), beta glucosidase (4) and glucosylceramidase (3). Interestingly, functional annotation by CUPP did not identify xylanolytic, chitinolytic or pectinolytic CAZymes. However, as the seagrass fibres (particularly of the *Zostera* genus) are known to be rich in both xylan and pectin (Davies et al. 2007) and the functional annotation by homology suggested xylanolytic and pectinolytic proteins (Data S1), we suggest that *Labyrinthula* SR_Ha_C could have such enzyme functionalities, but represented by enzymes too distantly related to the known CAZymes to be identified.

**Table 2.**
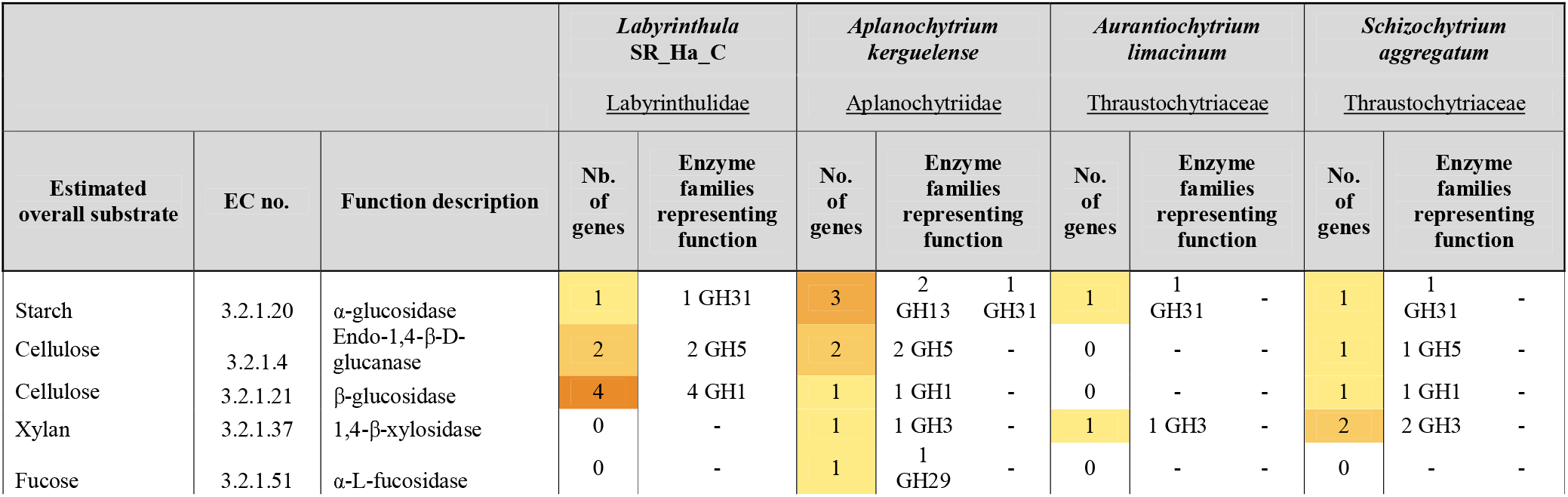
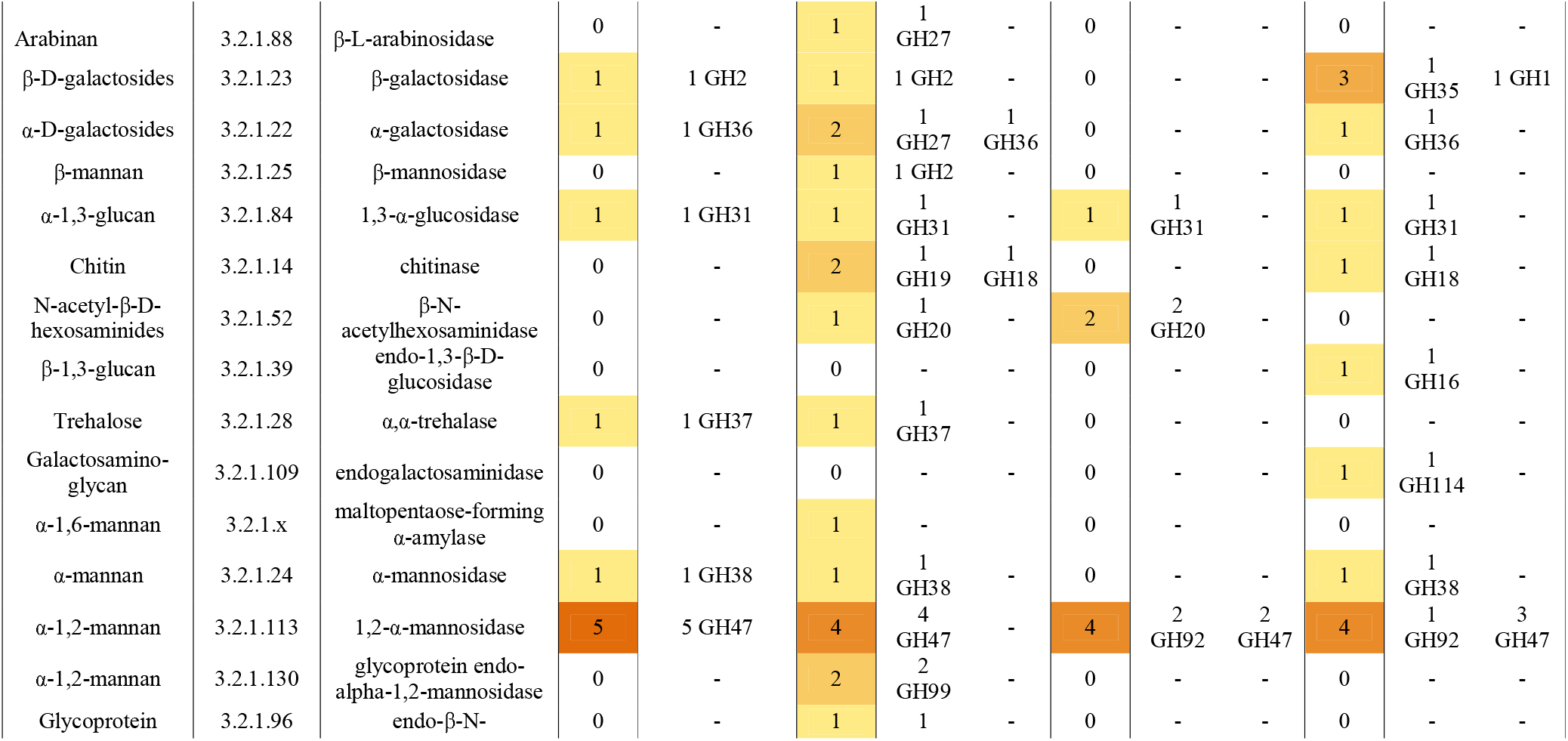
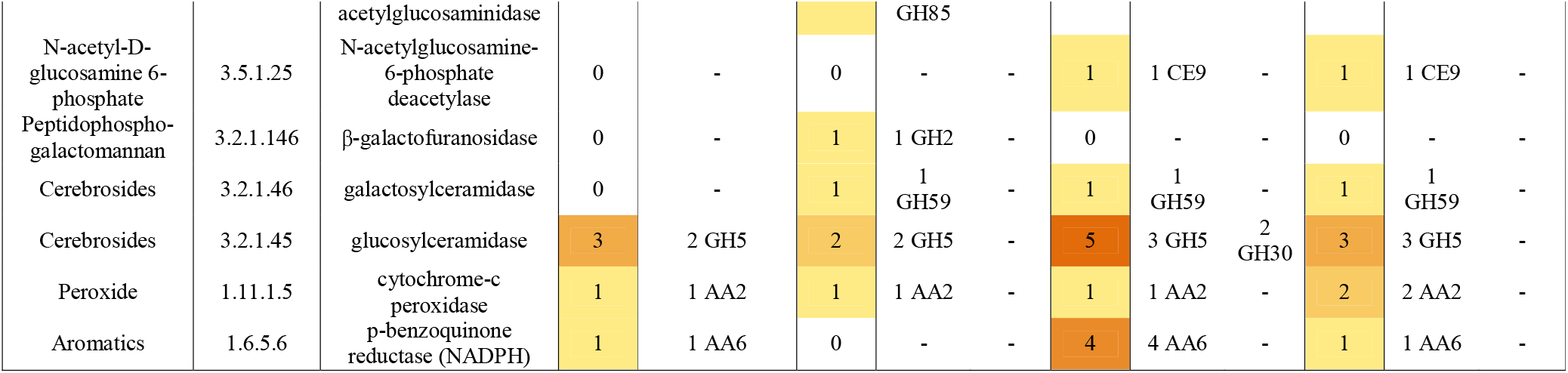
Comparative functional annotation of CAZymes in *Labyrinthula* sp. isolate SR_Ha_C as compared to three other Labyrinthulea. Each line of the table specifies one type of enzyme function, listing EC number, and estimated substrate. Note, the number of CAZymes functionally annotated for all these four species are as can be seen rather few. This is due to most of the secretome profile of the Labyrinthulea are too distantly related to known and characterised CAZymes to provide a basis for functional annotation. Further, as appears in the case of *Labyrinthula* SR_Ha_C, all enzyme functions are represented by only one type of enzyme family, while for the more diverse CAZyme profiles of the three other species, some enzyme functions are represented by two enzyme families. The Gene Number columns are given as heat maps. All CAZymes found by the CUPP analysis (both with predicted function and the CAZymes with no predicted function) are listed in Table S4.

The CUPP annotations of *Labyrinthula* SR_Ha_C were low (74) compared to the other three Labyrinthulea species: *A. kerguelens* (129), *S. aggregatum* (215), and *A. limacinum* (266; Tables 2, S4). Interestingly, the ratio of CAZymes, which could be predicted to function, divides the four species in two groups. Notably, the two Thraustochytrid species *A. limacinum* and *S. aggregatum* had 7% and 12 % of their CAZymes ascribed to function, respectively, while the two Labyrinthulidae species had 29% *(Labyrinthula* sp.) and 24% *(A. kerguelense*) ascribed. In short, among the four species studied, the richest CAZyme profile has also the highest level of uniqueness found among its enzyme portfolio (Tables 2, S4). Although we are working with a conservative predicted proteome, the rather low number of CAZymes found in *Labyrinthula* SR_Ha_C may be interpreted to reflect its ecology, e.g. being an opportunistic pathogen, it has a more specialised substrate/host affinity, as compared to the more saprotrophic lifestyles of the other three species. Further, it is attractive to interpret the more specialised enzyme profile of *Labyrinthula* SR_Ha_C. in the context of its highly specialised ectoplasmic net, a unique structure not only attaching the *Labyrinthula* colony to surfaces but also secreting digestive enzymes and nutritional uptake, similar to *S. aggregatum* (Iwata and Honda 2018).

*Labyrinthula* SR_Ha_C shared several Function:Family constellations with the other three Labyrinthulea species, that is, the four glycohydrolases, belonging to GH31 and GH38, are representing the same four EC functions in all species (Table 2). However significant discrepancies in Function:Family constellations are found as well: Enzyme function EC 3.2.1.113 predictive of α-mannan is represented by GH38 in *Labyrinthula* SR_Ha_C and *A. kerguelense*; while represented by two different proteins, GH47 and GH92 in the two Thraustochytriaceae species (*A. limacinum* and *S. aggregatum*). All in all, these observations reflect relatedness among the Labyrinthulea species, as well as indicating that they are only distantly related in digestive enzyme profiles, and likewise, substrate affinities.

### Evolutionary relationships within Bigyra and Stramenopile

We recovered a complete circular *Labyrinthula* SR_Ha_C mitogenome (accession number: MT267870) (Table 1, Figure S3). All genes were transcribed on the heavy strand, which has a guanine-rich base composition of 28.18% A, 8.26% C, 39.16% G and 24.40% T. In addition to the 13 protein-coding genes (PCG) typically found in animal mitogenomes, 10 other PCGs found in other heterotrophic protists were present in the *Labyrinthula* mitogenome (Wideman et al. 2020), namely *atp9, nad7, nad9, nad11, rp116, rps11-13, tatA* and *tatC*. At the time of this study, molecular sequences for *Labyrinthula* species were scant on the NCBI database, reporting only 572 nucleotide fragments, majority of which are ribosomal RNA and ITS sequences. Our complete annotated mitogenome showed an interesting strand asymmetry characteristic, containing a high guanine (G) to cytosine (C) ratio of 4.74:1 on its heavy strand versus an average of 1.22:1 in other Stramenopiles. Guanine-enriched sequences in other eukaryotes have been shown to adopt G-quadruplex (G4) stable structures and may play potential roles in the regulation of DNA replication, transcription and translation (Falabella et al. 2019). Using G4Hunter (Brázda et al. 2019), *Labyrinthula* SR_Ha_C’s predicted G4 frequency (15/1kbp) was 50- and 300-fold higher than predicted in *Schizochytrium* sp. TIO1101 and *Cafeteria roenbergensis*, respectively. It is difficult to infer that the cause of this high frequency due to the newness of this metric for non-human mitogenomes. However, as more mitogenomes are made available in the future, it will be of interest to see if this high frequency is observed in the mitogenomes of other more closely-related species and whether G4 richness is associated with protist evolutionary history or as potential regulators of virulence in pathogens (Harris and Merrick 2015).

In the reconstructed phylogeny of Stramenopiles, *Labyrinthula* and other protist species in the phylum, Bigyra formed a paraphyletic group at the base of Stramenopile, in addition to monophyletic clades for the other two phyla (Oomycota and Ochrophycota) (Figure 3; superalignment and phylogenetic tree is available in Data S2). Within Bigyra itself, *Labyrinthula* SR_Ha_C shared a strongly-supported sister relationship with two Thraustochytrida species (*Thraustochytrium aureum* and *Schizochytrium sp. TIO1101*), all three of which were taxonomically assigned to the class Labyrinthulea. This clade clustered with species from the other two major phyla, consistent with current taxonomic classifications and reports in other studies based on nuclear data (Tsui et al. 2009). However, this inference should be treated with caution since inadequate and uneven representation of branches in phylogenetic trees can introduce errors caused by long-branch attraction and false monophyly, which can lead to inaccurate phylogenetic relationships. It is therefore crucial to recognise the importance of strategic sampling to increase mitogenomic resources for under-studied taxonomic groups, thereby filling in the gaps in taxonomic representation to enable more comprehensive investigations into molecular diversity and evolutionary histories of protists (Wideman et al. 2020).

**Figure 3:**
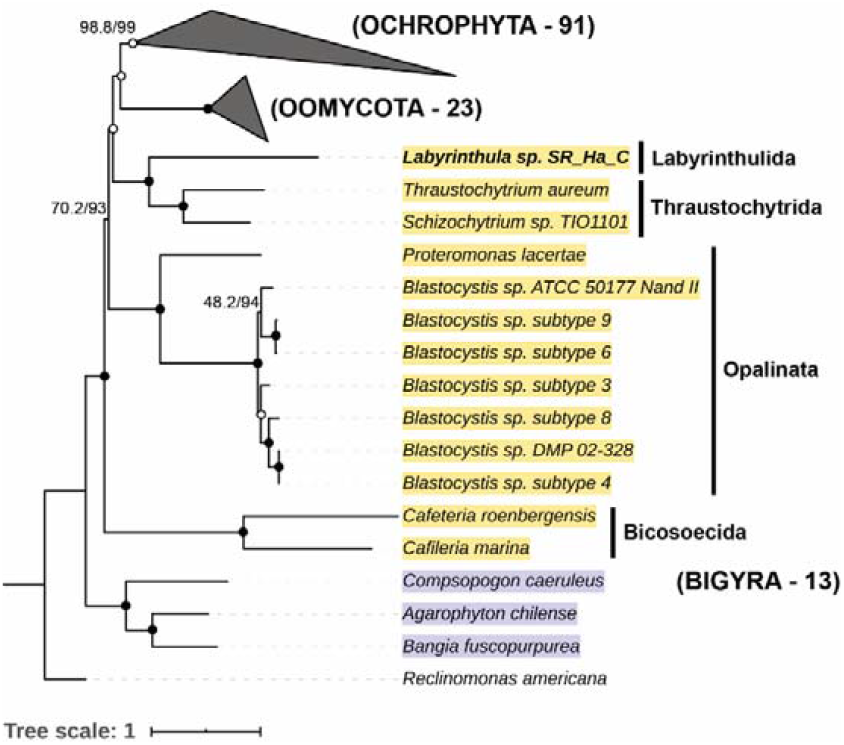
Evolutionary relationships of Labyrinthula sp. isolate SR_Ha_C and other species within Bigyra and Stremenopiles more generally. Phylogenetic tree based on 13 mitochondrial protein-coding genes (atp, cox, nad, cytb). This tree contains 127 Stramenopiles species/isolates in three phyla (Bigyra:13, Ochrophyta:91, Oomycota:23) and three Rhodophyta species as outgroup, rooted with Reclinomonas americana. Nodal support is indicated in the form of SH-aLRT / UFBoot values, with reliable clades having ≥80% and ≥95% values, respectively. Closed circles indicate maximum support (100/100) while open circles report less than maximum values but above the reliable threshold.

### Ecological and biotechnology implications of the *Labyrinthula SR_Ha_C* genome

This study reports the first assembled genome of a *Labyrinthula* species, and adds valuable genomic and transcriptomic resources to the limited molecular sequences available for the group Labyrinthulea (Song et al. 2018). The recovery of the complete mitochondrial genome makes this the first of its kind for the order Labyrinthulida, strongly supporting a monophyletic Labyrinthulida+Thraustochytrida clade, while providing insights into the evolutionary history of Stramenopile species. Our cross-phyla comparison of >100 Stramenopile genome assemblies unveiled possible caveats in current existing quality assessment tools that need to be carefully considered when assessing work for species from divergent and under-sampled taxonomic groups. Genome annotation of *Labyrinthula* SR_Ha_C opens comparisons to the recently published aquatic early lineage zoosporic fungi (Lange et al. 2019a). Further, the presented genome could act as a reference to propel new research directions for the *Labyrinthula* genus, such as population genetics (Lohan et al. 2020).

While our mining of the draft genome was not exhaustive, our initial investigations show promise in highlighting novel pathways to understanding *Labyrinthula* ecology and the SWD-pathosystem. The CAZymes and predicted CAZy function profiles provided insight into cell-wall breakdown. These pathways have been largely assumed through the classification of *Labyrinthula*’s saprobic lifestyle, but such evidence relating to this ecological function has most often been limited to their thraustochytrid cousins (Raghukumar and Damare 2011). Cell-wall and fibre breakdown not only has significant implications to marine carbon sequestration, but in conjunction with adhesion to the host, highlights mechanisms that may be key to the invasion of living seagrass tissue (Gibson et al. 2011, Iwata and Honda 2018). These results will hopefully provide a springboard to future research on *Labyrinthula*’s virulence or SWD assays using both host and pathogen gene responses.

Our study also highlights the industrial potential for *Labyrinthula*. Enzyme secretome profiles and specialised functional proteins of fungal and non-fungal zoosporic unicellular organisms, such as *Labyrinthula*, share many similarities with regard to enzyme functions, but also reflect different types of plant biomass affinities and degrading capabilities (Lange et al. 2019b). Secretome organisation, e.g. one enzyme function may be represented by several types of protein families or one type of protein family may include several enzyme functions, could reflect a strategy that is evolutionary optimised for robustness to varying conditions (salinity, temperature, pH, substrate availability). Notably, the uniqueness of the genome of *Labyrinthula* SR_Ha_C, combined with the very low number of sequenced genomes in this group makes the genome/transcriptome and function predictions described here a hotspot for future enzyme discovery activities. For example, the lignocellulose repertoire of plant pathogens is inspiring research on their use for biomass conversion to biofuels or higher value products (Gibson et al. 2011), while thraustochytrids are rich source bioproducts (Marchan et al. 2018). We expect that much more could be found from this highly diverse, un-exploited and under-studied group of microorganisms.

## Acknowledgments

This research utilised computational resources and services provided by the National Computational Infrastructure (NCI), which is supported by the Australian Government. We thank Samuel Lysy for his assistance in the lab. We would also like to thank Robert Ruge of Deakin University for assistance and use of the SIT HPC Cluster. Funding for this study was provided by the Mary Collins Trust and the Deakin Genomics Centre, Deakin University (Australia).

## Availability of data and material

The whole genome assembly was deposited into the NCBI/GenBank Whole Genome Sequence (WGS) database with the accession number JAALGZ000000000 and raw genomic reads into NCBI/Short Read Archive (SRA) with accession numbers SRR12496087 and SRR12496088, all of which are under the BioProject PRJNA607370. The mitochondrial genome was deposited into the NCBI/GenBank database with the accession number MT267870.

## Authors’ contributions

STT, SL, MT, LC, LL and FHG designed the research and direction of analyses. STT collected and isolated the sample. SL performed whole genome and transcriptome sequencing. MT assembled and annotated the genome and transcriptome, and carried out phylogenetic analysis. LC and MT performed comparative analyses of assemblies and proteomes of other protist genomes. LL, BP and MT analysed the peptide-based functional annotation of CAZymes. MT and STT led the writing of the manuscript. All authors edited and developed the manuscript.

