## Supplementary Methods for "First genome of *Labyrinthula*, an opportunistic seagrass pathogen, reveals novel insight into marine protist phylogeny, ecology and CAZyme cell-wall degradation"

### Summary supplementary and detailed Materials and Methods

### Note: Supplementary files other than Materials and Methods subject to request.

**Full Materials and Methods:** Below

**Figure S1**: Kmer-based estimations of haploid genome size and heterozygosity using GenomeScope (k=19, 21, 25).

**Figure S2**: Comparison of publicly-available genome assemblies in three phyla in Stramenopile (detailed version of Figure 1).

**Figure S3:** Circular mitogenome of *Labyrinthula* sp. isolate SR_Ha_C.

**Table S1**: List of species and corresponding GenBank accession numbers included in the genome tree.

**Table S2:** List of species and corresponding GenBank accession numbers included in the mitogenome tree.

**Table S3:** Detailed list of the carbohydrate-active enzyme genes and corresponding protein/transcript IDs in the *Labyrinthula* sp. isolate SR_Ha_C gene set (hyperlink tab).

**ESM 6: Table S4**: List of all CUPP annotated CAZymes from the four species of *Labyrinthulea*.

**Data S1:** Assembled transcriptome, gene model and sequences of predicted proteins/transcripts, and ortholog annotation output.

**Data S2:** Super-alignment of mitochondrial genes and phylogenetic tree in Newick format.

### Supplementary Materials and Methods

**Labyrinthula sp. *isolate description***

The *Labyrinthula* sp. culture used in this study was isolated from San Remo, Victoria, Australia in March 2016 (SR_Ha_C, 38.526305 S, 145.342938 E; Trevathan-Tackett et al. 2018). Briefly, the isolate, originally cultured from a green *Halophila* sp. leaf, was found to be highly pathogenic on both *Zostera* *muelleri* and *Heterozostera* *nigricaulis* (Trevathan-Tackett et al. 2018). Haplotype analysis revealed that it was closely related to another pathogenic *Labyrinthula* sp. isolate from Florida, USA and was within the putative pathogenic phylogenetic clade (Martin et al. 2016, Trevathan-Tackett et al. 2018). This isolate was chosen for sequencing, not only due to its virulence and close relatedness to *L. zosterae*, but the cells grow densely on top and within the culture media, which assisted in the extraction of a high concentration of high molecular weight genomic material. The isolate used for sequencing was cultured on seawater-serum agar plates at room temperature prior to sequencing, consisting of peptone, nutritional yeast, horse serum, glucose and Streptomycin-Penicillin antibiotics (Martin et al. 2009).

***gDNA extraction, library preparation and sequencing***

Genomic DNA for short-read Illumina sequencing was extracted from a 20 mg pellet of *Labyrinthula* SR_Ha_C culture (via scraping the top of the agar culture) with the ZymoBIOMICS Quick DNA kit according to the manufacturers’ instructions. The cell lysis step in the protocol was performed in the FastPrep-5G (MP Bio) for 40s at 5 m s^-1^ using a beating tube with a fine glass-bead matrix. The beat-beating solution was supplemented with 5 µl of RNaseA (40mg ml^-1^). The sample was incubated for 10 min at 50 °C after the addition of the lysis solution. Total RNA was prepared with 20 mg of culture using the Monarch Total RNA isolation kit according to the standard protocol for hard to lyse cells. A glass bead matrix was used for bead-beating lysis (30s at 5.5 m s^-1^). Integrity and purity of the DNA and RNA was confirmed using high-sensitivity TapeStation assays (Agilent), high sensitivity Qubit assays (Invitrogen) and by Maestronano (Maestrogen) UV microvolume spectrophotometer. The whole genome library was prepared with the Nextera Flex DNA library preparation kit (Illumina, San Diego) using 200 ng of purified genomic DNA as starting material. Short-read sequencing was performed on the Illumina MiSeq using 2x300 bp paired end sequencing. The short-read RNAseq library was prepared with 1ug of RNA with the Nugen mRNA ovation kit using the standard procedure. 15 PCR cycles were used to amplify the libraries. Sequencing was performed on the Illumina MiniSeq (2x150bp chemistry) generating almost 20 million pairs of reads.

Full-length transcriptome enriched libraries were prepared in triplicate from 1 µg of total RNA using Lexogen’s Teloprime full-length transcriptome kit, followed by 15 cycles of long-range PCR amplification. The full-length transcriptome libraries were pooled and prepared for long-read sequencing with sequencing kit SQK-LSK109 (ONT, UK). Sequencing was performed on one Flongle flow cell for 24 h and one MinION R9.4.1 flow cell (200 active starting pores, 200 mv voltage bias) for 24h. The Flongle (FLO) and MinION R9.4.1 flow cells were refuelled with a 50:50 mix of ultra-pure water and SQB buffer after 16h. Fast5 to fastq conversion was performed with guppy v 3.2. fast base calling algorithm (dna_R9.4.1_450bps_fast).

***Sequence processing and genome assembly***

Genomic raw data was generated in the form of short Illumina reads and long Nanopore reads. Illumina MiSeq reads were trimmed for quality and adapters with Trimmomatic v0.36 (*illuminaclip:2:30:10*, *avgqual:20*, *minlen:75*, *trailing:3*, *leading:3*) (Bolger et al. 2014), whereas adapters were trimmed from Nanopore MinION reads with Porechop v0.2.3 (Wick 2017).

Genome size for this *Labyrinthula* SR_Ha_C isolate was estimated based on the frequency of k-mers in short reads (Vurture et al. 2017). Illumina reads were first filtered to exclude reads of mitochondrial origin, an organelle that often exists in high copy numbers in a cell. This was done through alignment of reads to the mitogenome of *Labyrinthula* SR_Ha_C, which was also generated in this study (methods outlined below). Jellyfish v2.2.6 (Marçais and Kingsford 2011) was used to obtain distributions of 19-, 21- and 25-mer occurrences in pre-processed short reads, generating k-mer frequency histograms. These histograms were uploaded to the GenomeScope webserver (*max kmer coverage* disabled) (Vurture et al. 2017) for estimations of the haploid genome size, repetitive content and heterozygosity.

MaSuRCA v3.3.3 (Zimin et al. 2017) was used to perform a hybrid assembly of the *Labyrinthula sp.* SR_Ha_C genome, combining sequences from both short and long read data. Resulting contigs were further corrected and polished with three iterations of Racon v1.3.3 (Vaser et al. 2017) and Pilon v1.22 (Walker et al. 2014). Since bimodal k-mer distributions were observed from the GenomeScope analysis, which indicate relatively high levels of heterozygosity, the Purge Haplotigs pipeline (*-l 2 -m 12 -h 25*) (Roach et al. 2018) was used to remove haplotigs (i.e. artifactual contigs that exist in duplicates for highly heterozygous loci). One contig containing the *Labyrinthula* SR_Ha_C mitogenome was excluded from the final assembly. This assembly was assessed compared to that of other Stramenopiles based on assembly quality and completeness with Quast v5.0.2 (Mikheenko et al. 2018) and BUSCO v4.0.4 (Seppey et al. 2019), respectively, with the latter using single-copy genes from the Eukaryota and Stramenopiles datasets (*eukaryota_odb10*, *stramenopiles_odb10*). A list of species and corresponding GenBank accession numbers included in this comparison is available in Table S1.

Two checks were put into place to scan for possible bacterial contaminant contigs. The first was to search any contigs with first hits to bacterial sequences in NCBI’s non-redundant nucleotide database (*nt*). Secondly, RNAmmer v1.2 (Lagesen et al. 2007) was used to identify contigs containing bacterial ribosomal RNAs (*-S bac -m lsu,ssu,tsu*). In both searches, none of the contigs met these criteria and therefore all were retained.

***Mitogenome recovery and phylogenetic analysis***

A contig containing the complete *Labyrinthula* SR_Ha_C mitogenome was identified from the assembly through *blastn* homology with other protist mitogenomes and re-circularised. Preliminary annotation of the mitogenome was performed on the MFannot webserver (Beck and Lang 2010) and MITOS webserver (Bernt et al. 2013), followed by manual curation based on *blastp* homology to existing mitochondrial proteins in NCBI’s non-redundant (*nr*) database. The final annotated mitogenome was visualised with OGDRAW tool (Lohse et al. 2007).

Phylogenetic reconstruction was performed based on protein sequences from thirteen protein-coding genes typically found in eukaryotic mitogenomes (*atp6, atp8, cox1-3, nad1-6, nad4l, cytb*) (Boore 1999). These sequences were extracted from 127 mitogenomes representing 71 genera from the Stramenopile clade and 4 other mitogenomes as outgroup species (*Reclinomonas americana* and 3 Rhodophytes). Protein sequences of each gene were separately aligned with MAFFT (Katoh and Standley 2013), trimmed with trimAl (*-automated1*) (Capella-Gutiérrez et al. 2009) and subsequently concatenated with FASconCAT-G (Kück and Longo 2014) into one superalignment. This superalignment was used by IQ-TREE (*-m TESTMERGE*) (Nguyen et al. 2014) to find the best-fit partition model and to infer a phylogenetic tree based on maximum-likelihood, reporting nodal support in SH-aLRT (Guindon et al. 2010) and UFBoot (Minh et al. 2013) values. A clade with SH-aLRT ≥ 80% and UFBoot ≥ 95% is deemed reliable. The final phylogenetic tree was rooted with *Reclinomonas americana*, which has a mitogenome that most closely resembles the ancestral proto-mitochondrial genome (Lang et al. 1997). A list of species and corresponding GenBank accession numbers included in this tree is available in Table S2.

***Gene prediction and functional annotation***

Two types of “hints” were provided to increase accuracy in gene prediction for this non-model organism: (1) *Labyrinthula* SR_Ha_C transcript sequences, (2) known protein sequences. The first type consists of transcripts generated in this study by sequencing and assembling the transcriptome. Briefly, raw sequence reads generated by the Illumina MiniSeq were trimmed for poly-G with fastp v0.19.4 (Chen et al. 2018) and subsequently trimmed for quality and adapters with Trimmomatic v0.36 (Bolger et al. 2014). These reads were then assembled with Trinity v2.8.5 *(--SS_lib_type FR*) (Haas et al. 2013) to generate an Illumina-based transcriptome. Long cDNA reads sequenced on the Nanopore sequencer were used directly. The second type included known peptide sequences from 235 protists downloaded from Ensembl Protists (release 46) (Howe et al. 2019) and curated sequences from UniProtKB/Swiss-Prot (Consortium 2018).

The MAKER2 v2.31.10 annotation pipeline (Holt and Yandell 2011) was used to predict gene sequences from the assembled genome. A first iteration was run to identify a preliminary set of genes based on the aforementioned hints. These predicted genes were used to train two *ab initio* gene predictors, Augustus v3.2.3 (Stanke et al. 2006) and SNAP (Korf 2004), after which the resulting gene models were provided for a second MAKER2 iteration. Since genomes within the large protist clade are highly diverse and therefore contribute highly divergent gene sequences, the *keep_preds* option was activated in this second iteration to ensure all predictions are reported regardless of concordance to supporting hints. Instead, InterProScan v5.36-75.0 (Jones et al. 2014) was used to scan these predictions for the presence PFAM protein domains. Finally, any gene models that have physical evidence (AED < 1) and/or protein domain content were retained. For functional annotation, DIAMOND v0.9.24 (Buchfink et al. 2015) was used to find homology between predicted genes and publicly-available proteins in NCBI’s non-redundant protein database (*nr*).

Ortholog discovery within the kingdom Stramenopila (i.e. Chromista) was performed using protein sequences obtained from GenBank for two commonly-studied protists: 1. *Plasmodium falciparum* (BioProject: PRJNA148), 2. *Phytophthora sojae* (BioProject: PRJNA262907). Pairwise searches of protein sequence sets were performed with DIAMOND (no e-value cutoff) [28] and orthologs between *Labyrinthula* SR_Ha_C and each of the two species were identified through Reciprocal Best Hits (RBH).

***Identification of putative carbohydrate-active enzymes***

The dbCAN2 meta server (Zhang et al. 2018) was used to automate the annotation of carbohydrate-active enzymes (CAZymes) based on families of enzymes available in the CAZy database (Cantarel et al. 2008). Due to the divergence of *Labyrinthula* SR_Ha_C from other eukaryotes, the HMMER search option was selected (E-Value < 1e-15, coverage > 0.35) to search against representative hidden Markov models of signature domains of each CAZyme family within the five enzyme classes: 1. Glycoside Hydrolases (GH), 2. Glycosyl Transferases (GT), 3. Polysaccharide Lyases (PL), 4. Carbohydrate Esterases (CE), 5. Auxiliary Activities (AA). Counts of putative *Labyrinthula* SR_Ha_C CAZyme protein sequences were compared to existing data for 36 other Stramenopile species on the CAZy database.

Peptide based functional annotation of Labyrinthula CAZyme profile

The predicted amino acid sequences of *Labyrinthula* SR_Ha_C were analysed by the new non-alignment based method, Conserved Unique Peptide Patterns (CUPP) (Barrett and Lange 2019, Lange et al. 2019). CUPP automatically, with high precision and specificity, annotates proteins to families and subfamilies. If characterised members of the CUPP group query is already assigned to specific CAZyme function, CUPP assigns that function to the query protein. For the purpose of comparison, predicted proteins of three other members of Labyrinthulea (a.k.a. Labyrinthulomycetes) were analysed: *Aurantiochytrium limacinum* (ATCC MYA-1381), *Aurantiochytrium kerguelense* (PBS07) and *Schizochytrium aggregatum* (ATCC 28209); all three available from the Joint Genome Institute, DOE, US.
